# Histone methyltransferase DOT1L controls state-specific identity during B cell differentiation

**DOI:** 10.1101/826370

**Authors:** Muhammad Assad Aslam, Mir Farshid Alemdehy, Eliza Mari Kwesi-Maliepaard, Marieta Caganova, Iris N. Pardieck, Teun van den Brand, Fitriari Izzatunnisa Muhaimin, Tibor van Welsem, Iris de Rink, Ji-Ying Song, Elzo de Wit, Ramon Arens, Klaus Rajewsky, Heinz Jacobs, Fred van Leeuwen

**Affiliations:** Division of Tumor Biology and Immunology, Netherlands Cancer Institute, 1066CX Amsterdam, The Netherlands; Institute of Molecular Biology and Biotechnology, Bahauddin Zakariya University, 60800 Multan, Pakistan; Division of Gene Regulation, Netherlands Cancer Institute, 1066CX Amsterdam, The Netherlands; Max-Delbrück-Center for Molecular Medicine, 13125 Berlin, Germany; Immunohematology and Blood Transfusion, Leiden University Medical Center, 2333 ZA Leiden, The Netherlands; Division of Gene Regulation, Netherlands Cancer Institute, 1066CX Amsterdam, and Oncode Institute The Netherlands; Genome Core Facility, Netherlands Cancer Institute, 1066CX Amsterdam, The Netherlands; Division of Experimental Animal Pathology, Netherlands Cancer Institute, 1066CX Amsterdam, The Netherlands; Department of Medical Biology, Amsterdam UMC, location AMC, UvA, 1105 AZ Amsterdam, The Netherlands

**Keywords:** B cell differentiation, DOT1L, EZH2, Epigenetics, Germinal Center B cell, H3K79 methylation, Plasma cell

## Abstract

Differentiation of naïve peripheral B cells into terminally differentiated plasma cells is characterized by epigenetic alterations, yet the epigenetic mechanisms that control B cell fate remain unclear. Here we identified a central role for the histone H3K79 methyltransferase DOT1L in controlling B cell differentiation. Murine B cells lacking *Dot1L* failed to establish germinal centers (GC) and normal humoral immune responses *in vivo*. *In vitro*, activated B cells showed aberrant differentiation and prematurely acquired plasma cell features. Mechanistically, combined epigenomics and transcriptomics analysis revealed that DOT1L promotes expression of a pro-proliferative, pro-GC program. In addition, DOT1L supports the repression of an anti-proliferative, plasma cell differentiation program by maintaining expression of the H3K27 methyltransferase *Ezh2*, the catalytic component of Polycomb Repressor Complex 2 (PRC2). Our findings show that DOT1L is a central modulator of the core transcriptional and epigenetic landscape in B cells, establishing an epigenetic barrier that warrants B cell naivety and GC B cell differentiation.

## INTRODUCTION

B lymphocytes are key cellular components of the adaptive immune system and their functional deregulation is associated with immune deficiencies and autoimmunity^1, 2^. Several well-coordinated processes during B cell differentiation coincide with specific adaptations of the epigenome^3, 4, 5^. Given the critical contribution of B cells to the immune system, it is important to understand the molecular mechanisms underlying the epigenetic programming during their differentiation.

We previously identified DOT1L as a conserved epigenetic writer that catalyzes mono-, di-, or tri-methylation of lysine 79 of histone H3 (H3K79me)^6, 7^. DOT1L-mediated H3K79me resides on the nucleosome core surface away from histone tails ^8^ and is associated with active transcription. However, its function in gene regulation remains unclear^9, 10, 11^. DOT1L has gained wide attention as a specific drug target in the treatment of Mixed Lineage Leukemia (MLL) characterized by rearrangements of the *MLL* gene. Oncogenic MLL-fusion proteins recruit DOT1L, leading to hypermethylation of H3K79 and increased expression of MLL-target genes, thereby introducing a druggable dependency on DOT1L activity^12, 13, 14, 15, 16, 17, 18^. In addition, we recently observed a similar dependency in a mouse model of thymic lymphoma caused by loss of the histone deacetylase HDAC1^19^. While DOT1L is emerging as an appealing drug target, the role of DOT1L in gene regulation during normal lymphocyte development is not known. Analysis of publicly-available RNA-sequencing data shows that *Dot1L* expression is regulated during B cell development (see below).

An active humoral immune response is initiated by the expansion of clonally selected, antigen-primed B cells within secondary lymphoid organs. This results in the formation of a specific micro-environment, known as the germinal center (GC) ^20, 21^. Rapidly proliferating GC B cells pass through the process of somatic hypermutation that lays the molecular basis of antibody affinity maturation^22, 23, 24, 25, 26^. Ultimately, B cells selected on the basis of antibody affinity may differentiate either into memory B cells to establish long-term immunological memory or via a plasma blast stage into terminally differentiated, antibody secreting plasma cells. Development and functionality of B lymphocytes is associated with dynamic changes in the epigenetic landscape^27^. Recent studies indicate that specific alterations in B cell function and identity are intimately linked with well-established histone modifications such H3K4 trimethylation (H3K4me3) related with active gene promoters^28, 29^, and H3K27me3 associated with gene repression^30^. Furthermore, the H3K27 methyltransferase EZH2, the catalytic component of the Polycomb repressor complex 2 (PRC2) has been shown to have an essential role in establishing GC B cells^31^. Here we determined the role of the H3K79 methyltransferase DOT1L in normal murine B cell development by deleting *Dot1L* early in the B cell lineage and investigating specific dependencies of B lineage cells on DOT1L. Our findings show that DOT1L fine-tunes the core transcriptional and epigenetic landscape of B cells and in doing so establishes a critical epigenetic barrier coordinating the stepwise transitions towards terminally differentiated plasma cells.

## RESULTS

### Effective deletion of *Dot1L* in B-cell lineage cells

Given the DOT1L-dependencies in leukemia^13, 14, 32, 33^ and lymphoma^19^, we quantified the expression of *Dot1L* during normal B cell development using publicly available data^34^. We observed that *Dot1L* is transcriptionally regulated in B cell subsets and higher expressed in GC B cells (Fig. 1a). To determine the relevance of this regulation in controlling the development and differentiation of B lineage cells, we inactivated *Dot1L* during early B cell development by crossing the *Mb1-*Cre knock-in allele into a *Dot1L^fl/fl^* background. DOT1L is the sole enzyme responsible for H3K79me; knock-out of *Dot1L* has been shown to lead to complete loss of H3K79me^10, 19, 35, 36^. However, loss of H3K79 methylation requires dilution of modified histones by replication–dependent and –independent mechanisms of histone exchange^37, 38, 39^. *Mb1-*Cre was chosen because it leads to deletion of *Dot1L* at an early stage in B cell development that is followed by successive rounds of replication. This ensures complete loss of H3K79me in all subsequent B cell subsets. *Dot1L* was specifically and efficiently deleted in B cells, as confirmed by intracellular staining for H3K79me2 in proB, preB, immature B cells, and mature B cells. While some proB cells retained H3K79me2, preB cells and all stages beyond, lacked detectable levels of H3K79me2 (Fig. 1b). As a control, the methylation mark remained unchanged between DOT1L-proficient and - deficient mature T cells (Fig. 1b). We refer to *Mb1-*Cre^*+/−*^*;Dot1L^fl/fl^* and *Mb1-*Cre^*+/−*^*;Dot1L*^wt/wt^ B cells as *Dot1L* KO and WT cells, respectively. To study the impact of *Dot1L* ablation on the development of B cells in the bone marrow, we determined the cellularity of specific developmental subsets in the DOT1L-proficient and -deficient settings. Early ablation of *Dot1L* resulted in an overall 1.6-fold reduction of bone marrow B lineage cells. This reduction appeared to be caused primarily by an early differentiation block at the proB to preB cell stage; preB cell were reduced 1.8-fold in the *Dot1L* KO as compared to WT and proB cells were increased by 2.0-fold. In line with this partial developmental inhibition, the cellularity of all subsequent stages of development including immature B and mature B cells were significantly reduced in the bone marrow (Fig. 1c and Supplementary Fig. S1a). Reduced preB cells could be a result of impaired VDJ recombination in absence of DOT1L-dependent pro-recombinogenic H3K79me marks^40, 41^, but other causative factors cannot be excluded. Regardless of this partial developmental block, B cells could mature in the absence of DOT1L and H3K79 methylation, providing a system to study the role of this epigenetic mark in B cell differentiation.

**Figure 1:**
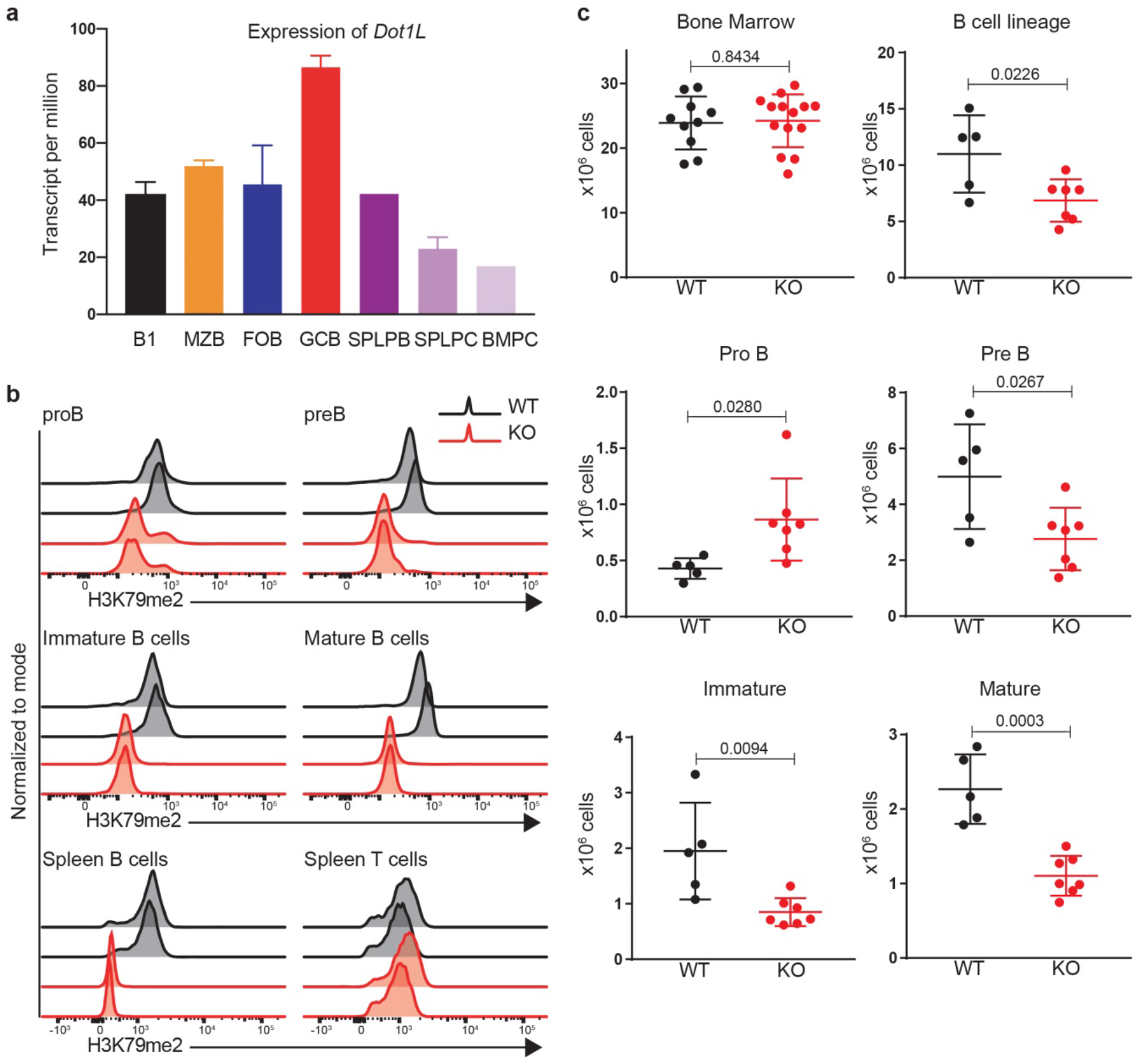
Expression levels, efficient deletion of Dot1L in B cells and its effect on cellularity of B lineage subsets in the bone marrow. a) Expression of *Dot1L* in different B-cell populations 34. B1: B1 cells, MZB: Marginal Zone B, FOB: Follicular B, GCB: Germinal Center B, SPLPB: spleen plasma blast, SPLPC: spleen plasma cells and Bone Marrow (BMPC: Bone marrow plasma cells). Expression is shown as transcript per million (TPM). Note, SPLPB and BMPC lack SD as replicates were lacking. b) Intracellular flow-cytometry staining for H3K79me2 in bone marrow B-cell subsets as well as splenic B and T cells from *Mb1-*Cre;*Dot1L^wt/wt^* (WT) and *Mb1-*Cre;*Dot1L^fl/fl^* (KO) mice. Results represent the data from two independent experiments. c) Absolute number of total nucleated cells from bone marrow B-cell subsets; p-value from Student’s t-test is indicated.

### Lack of DOT1L prohibits differentiation of germinal-center B cells

In the spleen of *Mb1-*Cre^*+/−*^*;Dot1L^fl/fl^* mice, B cell cellularity decreased 3.5-fold while T cell numbers remained unaffected, resulting in a 2.0-fold decreased overall cellularity (Fig. 2a-b). Among the various peripheral B cell subsets, the strongest reduction was found in marginal zone B cells and GC B cells to the extent that they were nearly absent (Fig. 2c-i and Supplementary Fig. S1b). The reduction of GC B cells was of particular interest given the highest levels of *Dot1L* mRNA expression in this subset (Fig. 1a and 2i), and strongly suggested that the formation of germinal centers critically depends on DOT1L. Indeed, *in situ* histological analyses also revealed the absence of GCs in the spleen of *Dot1L*-KO mice (Fig. 2j and 2k). Similarly, lack of DOT1L resulted in a marked reduction of GC B cells in Peyer’s patches (Supplementary Fig. S2).

**Figure 2:**
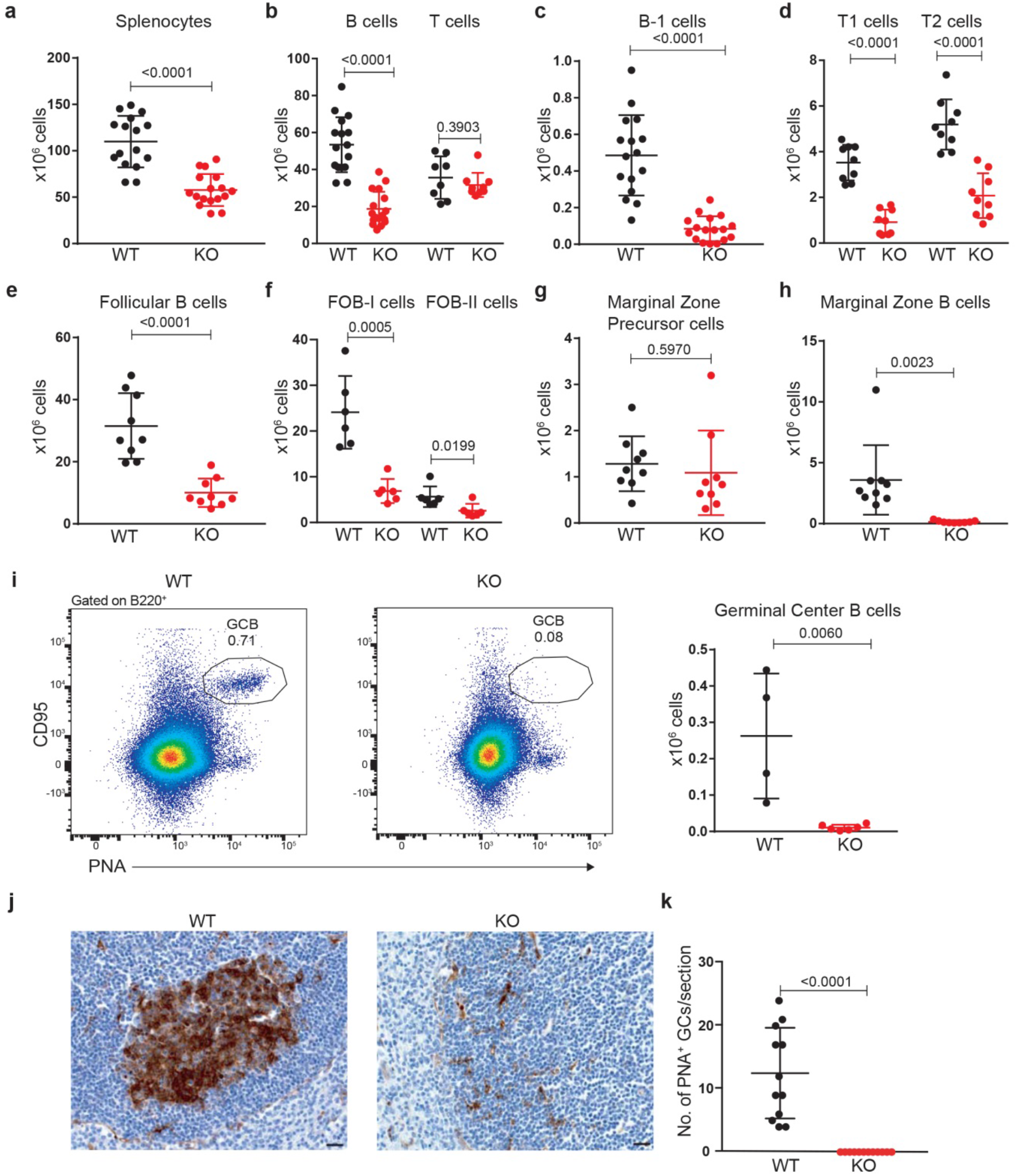
Dot1L ablation affects the cellularity of mature peripheral B cell populations with a strong reduction in Marginal Zone and Germinal Center B cells. (a-h) Absolute number of total nucleated splenocytes, splenic B-and T cells, and indicated mature B cell subsets in *Mb1*-Cre;*Dot1L^wt/wt^* (WT) and *Mb1*-Cre;*Dot1L^fl/fl^* (KO) mice; p-value from Student’s t-test is indicated. Error bars indicate mean +SD. (i) Number of germinal center B cells (PNA^high^, CD95^+^) and flow cytometric analysis of GC B cells from the spleen of unchallenged WT and KO mice; p-value from t-test is indicated. Error bars indicate mean ± SD (j and k). The quantification and identification of germinal centers lectin histochemistry of Peanut agglutinin (PNA) in spleens from WT and KO mice. The scale bar: 20 μm. Error bars indicate mean with ± SD.

### Dot1L deletion impairs proliferation *in vitro*

Following B-cell priming, B cells undergo class switch recombination (CSR) and form GCs. CSR is an important feature of humoral immunity and is dependent on B cell activation and initiation of proliferation. Given the critical role of DOT1L in GC B cell differentiation, we here examined the CSR potential and proliferative capacity of naïve splenic B cells stimulated *in vitro* with LPS alone or in combination with IL-4, to induce switching to IgG3 or IgG1, respectively. Compared to *Dot1L*-proficient B cells, the lack of *Dot1L* was associated with a reduced frequency of class-switched IgG3 and IgG1 cells (Fig. 3a-b, and Supplementary Fig. S3a-b). Similar observations were made using the T-cell dependent mimetic anti-CD40 and IL-4 as stimuli (Fig. 3c and Supplementary Fig. S3c). Mechanistically, this reduction in CSR might relate to impaired proliferation, since proliferation is a prerequisite for CSR ^42^ (e.g. see WT in Fig. 3d-e), or a defect in the CSR machinery. We found that the proliferative response of *Dot1L*-KO B cells was strongly impaired upon CD40+IL-4 *in-vitro* stimulation, as revealed by tracing the dilution of a fluorescent label (Fig. 3d). The reduced CSR potential is most likely caused by the impaired proliferation as the frequency of switched cells among equally-proliferating populations was not different between KO and WT (Fig. 3e). Furthermore, RNA-Seq analysis of KO and WT B cells activated with LPS and IL-4 argued against a potential failure of CSR in *Dot1L-*KO B cells, since components of the CSR machinery were expressed in KO cells (Supplementary Table S1). Finally, *in-vitro* activation of naïve B cells with LPS and IL-4 led to a higher proportion of dead cells (Zombie NIR positive) in *Dot1*L-KO compared to WT. In addition, the viable (Zombie NIR negative) subset in activated KO B cells consistently showed higher levels of active caspase 3, a marker of apoptosis. These insights indicated that the viability of *Dot1L*-deficient B cells is severely compromised (Supplementary Fig. S3d-f). The virtual absence of GC B cells, the impaired proliferative response, and the increased cell death of *Dot1L*-deficient B cells implicated a severe defect of *Mb1-*Cre^*+/−*^*;Dot1L^fl/fl^* mice in establishing effective humoral immunity.

**Figure 3:**
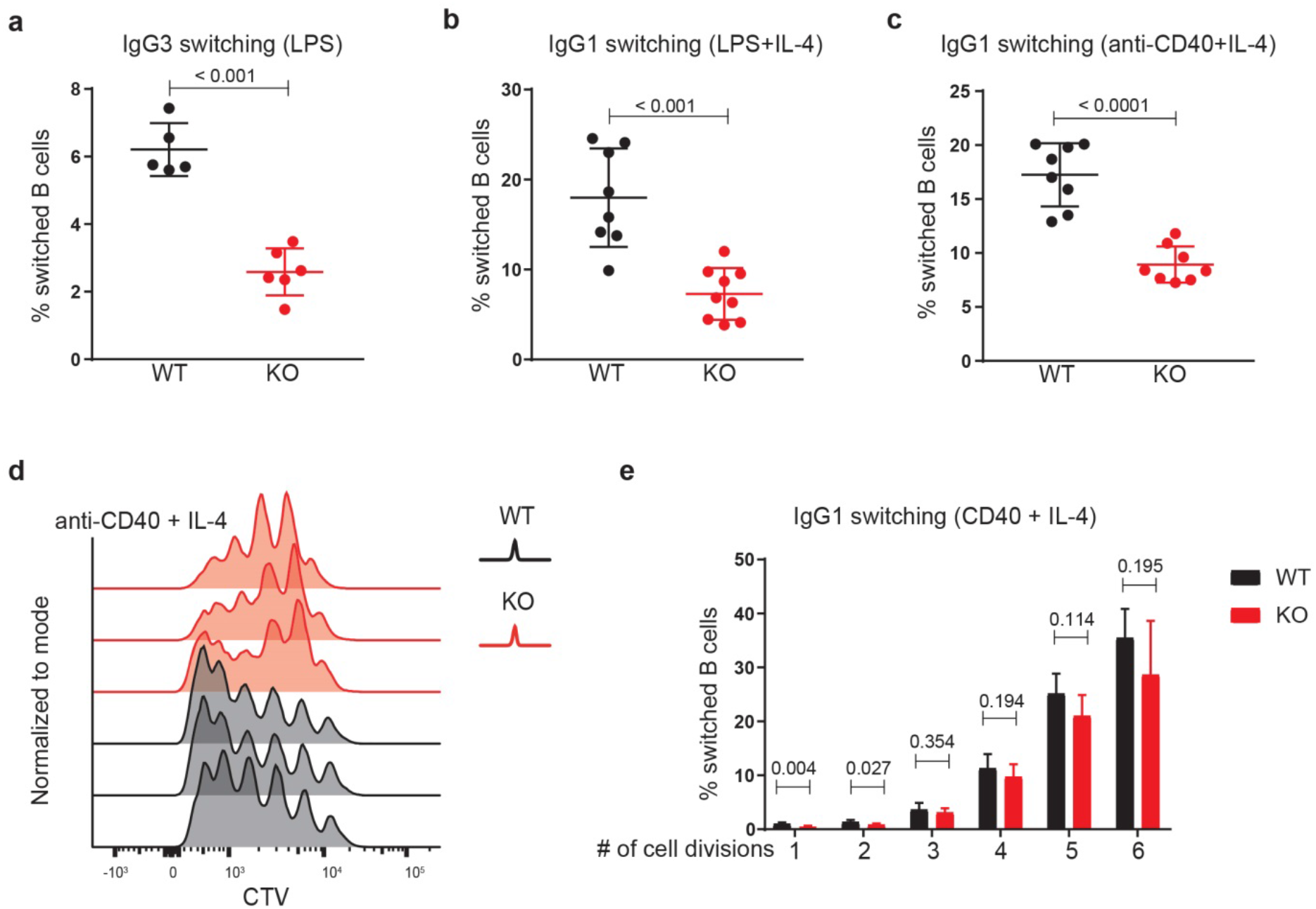
DOT1L is essential for efficient CSR and proliferation. (a-b) Analysis of DOT1L-proficient (WT) and DOT1L-deficient (KO) naïve B cells activated for four days with LPS alone (IgG3 switching) or with LPS and IL-4 (IgG1 switching); p-value from Student’s t-test is indicated. Error bars indicate mean ±SD. c) Analysis of DOT1L-proficient (WT) and -deficient (KO) B cells, activated for four days with anti-CD40 and IL-4 (IgG1 switching) B cells. The graph indicates the statistical analysis (Student’s t test) of the percentages of B cells that switched to IgG1 in WT and KO; p-value from t-test is indicated. Error bars indicate mean ±SD. d) Naïve B cells were labeled with cell trace violet (CTV) and stimulated for four days with anti-CD40 and IL-4. CTV dilution was measured by flow cytometry. e) Percentage of IgG1 switched cells per generation of proliferating cells. p-value greater than 0.05 is considered as non-significant. Results from three independent WT and KO mice are depicted.

### *Dot1L*-deficient B cells fail to mount efficient immune responses *in vivo* and acquire plasma cell features *in vitro*

In unchallenged *Dot1L*-KO mice, serum IgM titers appeared normal but IgG1 titers were decreased, while IgG2b, IgG3, and IgA titers were not significantly decreased (Supplementary Fig. S4a-e). To determine the immune responsiveness of *Dot1L*-deficient B cells, *Mb1-*Cre^*+/−*^*;Dot1L^fl/fl^* and *Mb1-*Cre^*+/−*^*;Dot1L^wt/wt^* mice were challenged with an acute lymphocytic choriomeningitis virus (LCMV) infection. During the LCMV infection, *Dot1L*-deficient B cells failed to establish GC B cells and to generate plasma cells (Fig. 4a-b). In accordance with this, these mice also failed to mount normal IgM serum titers against LCMV (Fig. 4c). The failure to establish GCs in response to LCMV is also in line with the very low LCMV-specific IgG serum titers (Fig. 4c). The inability of *Mb1-*Cre^*+/−*^*;Dot1L^fl/fl^* mice to mount efficient antibody responses to T-cell dependent antigen was confirmed using 4-hydroxy-3-nitrophenylacetyl conjugated to chicken gamma globulin (NP-CGG) in alum as immunogen (Fig. 4d). Together, these data suggest that DOT1L is essential for establishing a normal humoral immune response. Upon *in-vitro* activation of naïve *Dot1L*-KO B cells with LPS and IL-4, a significantly increased frequency of cells expressing the plasma-cell marker CD138 ^43, 44^ was observed (Supplementary Fig. S4f). This observation was supported by the increased proportion of cells expressing the pan-plasma cell transcription factor BLIMP1 ^45^ (Supplementary Fig. S4g) in *Dot1L-*KO versus WT. However, these cells failed to downregulate B220 (Supplementary Fig. S4h). Downregulation of B220 is considered as a hallmark of post-mitotic plasma cells ^46^. In addition, *in-vitro* activated *Dot1L*-KO B cells failed to differentiate into CD19-high activated B cell blasts or CD19-negative plasma cells, but were stuck at a transitional state, expressing CD19 at intermediate levels (Supplementary Fig. S4i). Together, these phenotypes upon *in-vitro* activation suggest that in the absence of DOT1L-mediated H3K79 methylation, naïve B cells skip the GC stage and prematurely start to gain some plasma cell features but do not accomplish complete differentiation program.

**Figure 4:**
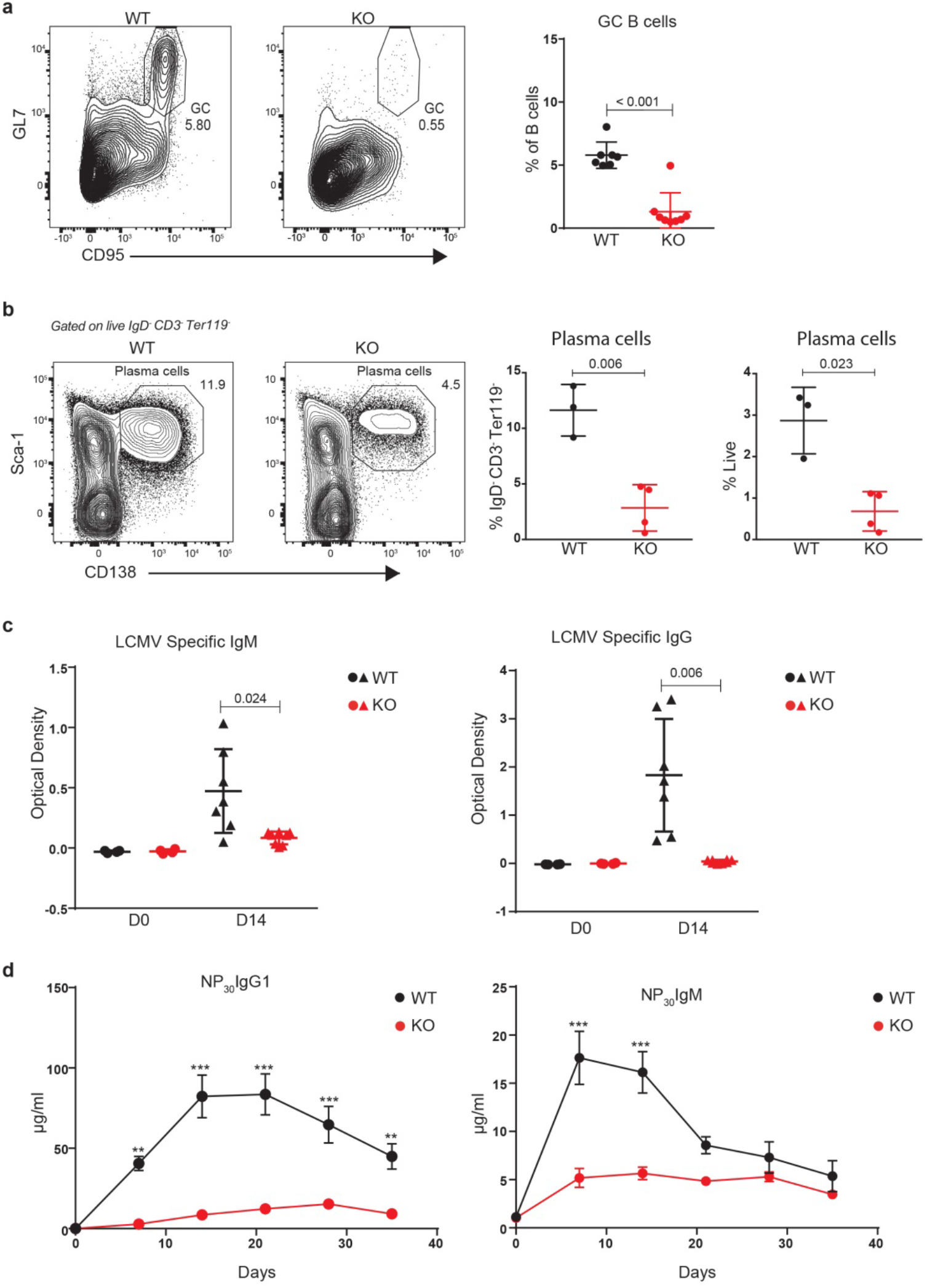
DOT1L-deficient B cells fail to mount an efficient immune response but show signs of plasma-cell differentiation upon challenge. a) Representative flow cytometry plots of splenic GC B cells 14 days after LCMV Armstrong infection in WT and *Dot1L*-KO mice and their statistical analyses. Error bars indicate mean with ± SD. b) Representative flow cytometry plots of plasma cells B from the spleen at 14 days after LCMV infection and their statistical analyses. Error bars indicate mean with ± SD. c) LCMV-specific IgM and IgG titers in the serum of WT and KO mice before (D0) and 14 days after (D14) infection; number indicates p-value of a Student’s t-test. Error bars indicate mean with ± SD. d) WT and KO mice were challenged with the model antigen 4-hydroxy-3-nitrophenylacetyl conjugated to chicken gamma globulin (NP-CGG). NP-specific IgG1 and IgM titers were quantified by ELISA from the serum isolated at the indicated days following immune challenge. Error bars indicate mean ± standard error of mean (SEM).

### DOT1L supports a pro-proliferative, MYC-high GC stage and prohibits premature differentiation towards plasma cells *in vitro*

To unravel the underlying molecular mechanisms that prohibit *Dot1L*-KO GC B cell differentiation and stimulate partial differentiation towards plasma cells we performed RNA-Seq analyses of *in vitro* activated (LPS and IL-4) B cells under *Dot1L*-proficient and -deficient conditions. Among the differentially-expressed genes, the genes encoding the pro-GC transcription factor BACH2 and the pro-proliferative transcription factor MYC^25, 47, 48^ were H3K79me2 methylated in WT cells and transcriptionally downregulated in activated DOT1L-deficient B cells (Supplementary Fig. S5a-d). In agreement with this, *Prdm1*, repressed by BACH2 (and H3K79me2-methylated and expressed in activated B cells) and encoding the pan-plasma cell transcription factor BLIMP1^45, 49^, was found upregulated (Fig. 5b and Supplementary. Figs. S5c-d). These transcriptional changes are in line with the observed absence of GC B cells and increased formation of cells with plasma cell features in KO cells. In addition, analysis of a published plasma-cell gene signature ^34^, revealed that the transcriptome of activated *Dot1L*-deficient B cells was indeed strongly enriched for plasma-cell associated transcripts (Fig. 5c). Simultaneously, MYC-target gene transcripts were strongly reduced (Fig. 5d). Together, these observations indicate that DOT1L-mediated H3K79 methylation licenses a transient entrance into a pro-proliferative, MYC-high GC stage and is required to prevent premature differentiation of activated B cells towards non-proliferative terminally differentiated plasma cells. Transcriptomic data from *ex-vivo* isolated naïve *Dot1L*-deficient B cells also revealed enrichment of some plasma-cell associated genes, suggesting that antigen-inexperienced B cells are already prematurely differentiated in the absence of DOT1L (Supplementary Fig. S5e). However, upon activation, expression of *Irf4*, a factor essential for plasma cell differentiation^50^, was not differential between WT and KO. This dichotomy, amongst other factors, likely contributes to the failure to adopt an identity of terminally differentiated plasma cells (Supplementary Fig. S5f). Together, our results indicate that DOT1L modulates the expression of several key transcriptional regulators essential in controlling stepwise B cell differentiation.

**Figure 5:**
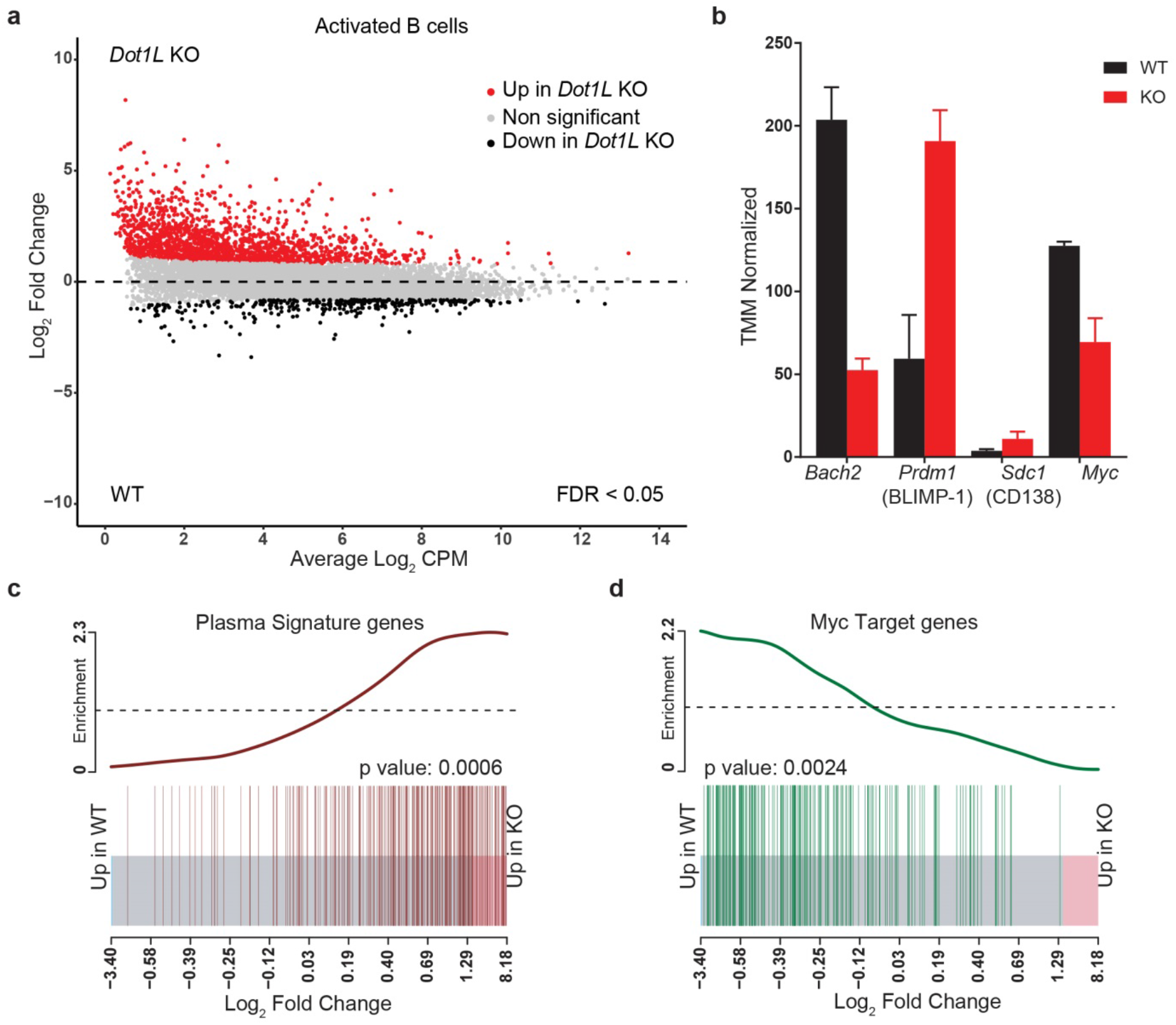
Transcriptome analysis of *in-vitro* activated B cells shows accelerated plasma-cell differentiation and compromised activation of MYC-target genes in the absence of DOT1L. a) MA-Plot of normalized RNA-Seq data showing differentially expressed genes (FDR < 0.05) between *Dot1L*-KO and WT B cells after two days of *in-vitro* activation with LPS+IL-4. b) Differential expression of *Bach2*, *Prdm1* (encoding BLIMP-1), *Cd138* (a plasma-cell marker) and *Myc* transcripts are indicated as CPM after TMM normalization from WT and KO. Error bars indicate mean ± SD. c) BARCODE plot showing the enrichment of plasma-cell signature genes in KO as compared to WT activated B cells. d) BARCODE plot showing the enrichment of MYC-target genes in WT as compared to KO activated B cells. p values showing the statistical significance of enrichment of gene set calculated via FRY test are indicated.

### DOT1L-mediated H3K79 methylation is associated with gene activity in B cells

To link the phenotypes of loss of DOT1L to its role as an epigenetic regulator in B cells, we generated genome-wide maps of DOT1L-mediated H3K79me2 in naïve and activated WT B cells by ChIP-Seq. We analyzed the H3K79me2 mark because it is known to mark the region downstream of the transcriptional start site of transcribed genes and positively correlate with gene activity^9, 10, 51, 52^. While H3K79me1 shows the same trends, it has a broader distribution, and H3K79me3 is detectable only at limited levels^9, 19, 52^. Here we analyzed how the gene-expression changes in *Dot1L*-KO versus WT B cells related to genes being marked by H3K79me2 in WT cells.

Both in naïve and activated B cells more than 83 percent of the differentially expressed genes was found upregulated in the absence of DOT1L; only a small subset was downregulated (Fig. 5a and Supplementary Fig. S5g). The upregulation was biased towards more lowly expressed genes. The observed upregulation of genes in *Dot1L*-KO B cells was unexpected given the fact that H3K79me2 generally correlates with transcriptional activity^9, 11, 17, 51, 53, 54, 55, 56^. However, repressive functions of DOT1L have been proposed as well^57, 58, 59, 60^. Comparing H3K79me2 ChIP values in WT cells with the gene expression changes caused by loss of DOT1L revealed that the genes upregulated in *Dot1L*-KO B cells were mostly hypomethylated in WT cells, indicating that they are likely indirectly affected by the loss of DOT1L (Fig. 6a and 6b). In contrast, genes downregulated in *Dot1L*-KO B cells were generally highly expressed and H3K79 methylated in WT B cells, indicating that this gene set harbors the genes directly dependent on DOT1L. These findings show that DOT1L-mediated H3K79me2 is a mark of many active genes in B cells, but suggest that only a small fraction of these genes requires H3K79me2 for maintenance of gene expression, since only a subset of the active genes was downregulated in *Dot1L* KO.

**Figure 6:**
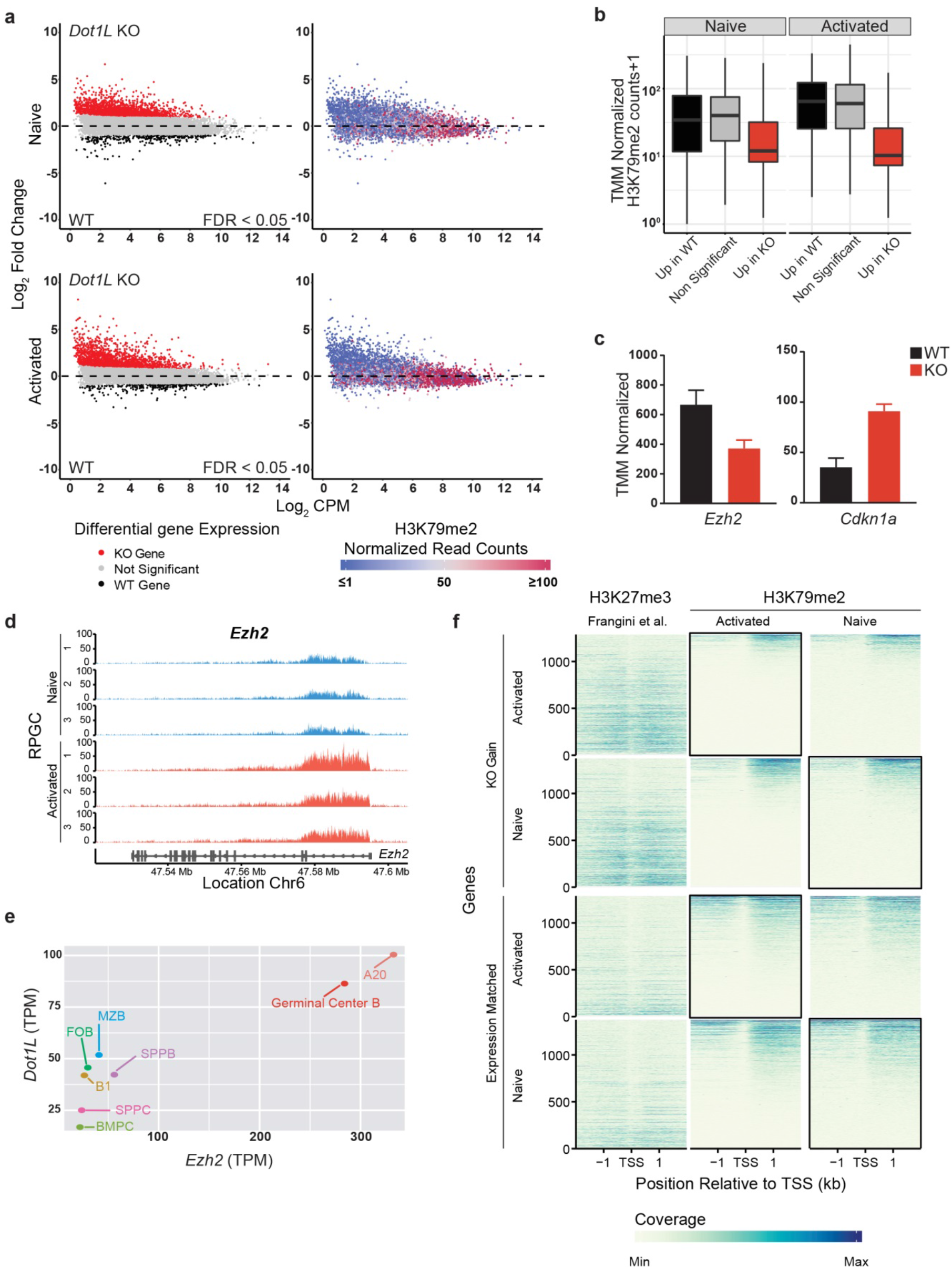
DOT1L-mediated H3K79 methylation is associated with gene activity in B cells and indirectly promotes repression of PRC2 target genes. a) Integrative analyses of differentially expressed transcripts (FDR < 0.05) from activated and naïve Dot1L-deficient B cells (left panel) with H3K79me2 ChIP values from WT activated and naïve B cells (right panel). b) The distribution of mean H3K79me2 amongst different gene sets from activated and naïve B cells as depicted by box plots. c) Differential expression of *Ezh2* and *Cdkn1a* is depicted as CPM after TMM normalization from WT and *Dot1L*-KO activated B cells. Error bars indicate mean ± SD. d) H3K79me2 methylation at the *Ezh2* locus from WT activated and naïve B cells, as determined by reads per genomic content (RPGC); three independent replicates are shown. e) Scatter plot showing the correlation between expression of *Ezh2* and *Dot1L* as depicted in TPM in different mature B-cell subsets; B1: B1 cells, MZB: Marginal Zone B, FOB: Follicular B, GCB: Germinal Center B, SPLPB: spleen plasma blast, SPLPC: spleen plasma cells) and Bone Marrow (BMPC: Bone marrow plasma cells) A20: Germinal Center like cell lymphoma cell line. f) Coverage plot of H3K27me3 from naïve B cells 67 and H3K79me2 (from WT naïve and activated B cells) flanking four kb around transcriptional start sites (TSS) for genes upregulated in KO (KO Gain) or non-differential Expression Matched genes is shown. Coverage was calculated as reads per genomic content cutoff at the 0.995th quantile and rescaled to a maximum of 1.

### DOT1L supports repression of PRC2 target genes

The large number of genes found upregulated in *Dot1L*-KO B cells indicated that DOT1L positively regulates the expression of a transcriptional repressor whose target genes are derepressed in absence of DOT1L. To identify such candidate repressors, we investigated the relatively small fraction of genes downregulated in naïve and activated *Dot1L*-KO B cells. Unbiased identification of upstream transcriptional regulators by Ingenuity Pathway analyses (IPA) pointed towards potential regulators that were differentially upregulated in WT B cells (Tables 1 and 2). To further narrow down the list, we filtered for genes that are (i) H3K79-dimethylated by DOT1L, (ii) encode transcriptional repressors, and (iii) play a role in B cell differentiation/proliferation and GC formation. This led to the identification of the histone H3K27 methyltransferase EZH2 as a candidate factor (Table 1 and 2). We verified that *Ezh2* expression was reduced in *Dot1L*-KO B cells (Fig. 6c) and that the gene is H3K79-dimethylated in WT cells (Fig. 6d), indicating that the expression of *Ezh2* is directly promoted by DOT1L activity (Fig. 6c and 6d). Further supporting this notion, the expression of *Ezh2* and *Dot1L* was found co-regulated in B cell subsets (Fig. 6e).

**Table 1:** Ingenuity Pathway Analysis of upstream transcriptional regulators that are differentially expressed higher in WT than DOT1L knock-out naive B cells (FDR < 0.05)

| Upstream Transcriptional Regulator | Expression Log Ratio (KO/WT) | Activation z-score | p-value of overlap | Ensemble gene id | TMM normalized H3K79me2 |
| --- | --- | --- | --- | --- | --- |
| *HIF1A | -1.534 | 5.138 | 3.22E-10 | ENSMUSG00000021109 | 8.44 |
| EZH2 | -1.195 | 0.27 | 0.000133 | ENSMUSG00000029687 | 33.46 |
| REL | -1.146 | 1.685 | 9.25E-05 | ENSMUSG00000020275 | 41.44 |
| TAF4B | -1.042 |  | 0.0396 | ENSMUSG00000054321 | 23.16 |
| IRF4 | -1.038 | -3.202 | 0.00193 | ENSMUSG00000021356 | 120.62 |
| MED13 | -0.978 | -2.143 | 0.0067 | ENSMUSG00000034297 | 153.69 |
| *SIRT1 | -0.973 | -3.106 | 7.11E-09 | ENSMUSG00000020063 | 7.74 |
| PURA | -0.964 | 1.067 | 0.00223 | ENSMUSG00000043991 | 47.75 |
| EBF1 | -0.95 | 2.612 | 7.55E-08 | ENSMUSG00000057098 | 112.63 |

\* Genes with TMM normalized H3K79me2 score less than 10 were not considered.

**Table 2:** Ingenuity Pathway Analysis of upstream transcriptional regulators that are differentially expressed higher in WT than DOT1L knock-out activated B cells (LPS+IL-4, Day 2; FDR < 0.05)

| Upstream Transcriptional Regulator | Expression Log Ratio (KO/WT) | Activation z-score | p-value of overlap | Ensemble gene id | TMM normalized H3K79me2 |
| --- | --- | --- | --- | --- | --- |
| BACH2 | -1.956 | -3.367 | 0.00186 | ENSMUSG00000040270 | 66.81 |
| *EGR1 | -1.678 | 3.22 | 6.76E-15 | ENSMUSG00000038418 | 10.41 |
| TAF4B | -1.436 |  | 0.00102 | ENSMUSG00000054321 | 53.19 |
| PURA | -1.392 |  | 0.00633 | ENSMUSG00000043991 | 49.35 |
| *HIF1A | -1.269 | 4.754 | 7.83E-10 | ENSMUSG00000021109 | 6.13 |
| CREBZF | -1.233 | -1.994 | 0.0409 | ENSMUSG00000051451 | 206.73 |
| SMAD2 | -1.178 | 2.439 | 0.0061 | ENSMUSG00000024563 | 94.23 |
| EBF1 | -1.15 | 1.207 | 0.000138 | ENSMUSG00000057098 | 155.78 |
| HHEX | -1.059 | -0.538 | 0.00453 | ENSMUSG00000024986 | 41.60 |
| *EGR2 | -1.018 | 1.566 | 1.16E-06 | ENSMUSG00000037868 | 8.55 |
| PDLIM1 | -0.967 |  | 0.0383 | ENSMUSG00000055044 | 16.33 |
| SATB1 | -0.962 | -1.998 | 4.34E-07 | ENSMUSG00000023927 | 116.85 |
| CREB1 | -0.949 | 3.751 | 6.80E-13 | ENSMUSG00000025958 | 92.51 |
| PAX5 | -0.917 | -1.807 | 0.00318 | ENSMUSG00000014030 | 330.58 |
| KLF2 | -0.912 | 1.144 | 1.98E-15 | ENSMUSG00000055148 | 148.02 |
| NPM1 | -0.881 | -0.509 | 0.00309 | ENSMUSG00000057113 | 177.98 |
| MYC | -0.878 | -3.848 | 2.74E-14 | ENSMUSG00000022346 | 163.26 |
| REL | -0.866 | 1.473 | 1.58E-10 | ENSMUSG00000020275 | 102.56 |
| EZH2 | -0.846 | -0.678 | 2.11E-06 | ENSMUSG00000029687 | 135.73 |
| *CBFB | -0.814 | 0.538 | 2.45E-05 | ENSMUSG00000031885 | 6.84 |
| NFKB1 | -0.813 | 2.31 | 6.65E-12 | ENSMUSG00000028163 | 176.38 |

\* Genes with TMM normalized H3K79me2 score less than 10 were not considered.

We next investigated the physiological relevance of the connection between DOT1L and EZH2. First, *Ezh2*-KO and *Dot1L*-KO B cells have overlapping phenotypes^31, 61, 62, 63, 64, 85^. Second, we took advantage of publicly available RNA-Seq data^62^ from *Ezh2*-KO plasma cells to identify genes that require EZH2 for their repression (Supplementary Fig. S6a). Using this as a signature of EZH2-dependent genes, we found that many of the genes de-repressed in *Ezh2*-KO cells were also found de-repressed in activated *Dot1L*-KO B cells (Supplementary Fig. S6b). Many of these genes were found de-repressed also in naïve *Dot1L*-KO B cells (Supplementary Fig. S6b). As an independent validation, we evaluated the expression of a known PRC2-target gene *Cdkn1a* (*p21*)^63, 65, 66^, and found that it was upregulated in activated *Dot1L*-KO B cells (Fig. 6c). Third, we analyzed the level of H3K27me3 and H3K79me2 in the set of genes that was de-repressed in *Dot1L* KO using our H3K79me2 ChIP-Seq data and publicly available H3K27me3 ChIP-Seq data from naïve B cells^67^. The analysis revealed that this gene set was enriched for H3K27me3 in WT cells compared to expression-matched non-differentially expressed genes (Fig. 6f). Together, these findings suggest that in B cells, DOT1L supports the repression of PRC2 target genes, thus uncovering a previously unknown connection between two conserved histone methyltransferases associated with activation and repression, respectively. Together, our findings place H3K79 methylation by DOT1L at the heart of maintaining epigenetic identity of B cells, by orchestrating the activity of central transcriptional and epigenetic regulators such as BACH2, MYC, EZH2, and their target genes.

## DISCUSSION

Given the critical contribution of B cells to the immune system, it is important to understand the molecular mechanisms underlying the epigenetic programming during their differentiation. Taking advantage of a B-cell specific murine knock-out model, we observed that the H3K79 methyltransferase DOT1L has a central role in B cell physiology. Among mature B cells, GC B cells express the highest levels of *Dot1L* and they were strongly reduced in *Dot1L*-KO mice. Indeed, GC B cell differentiation was found to be critically dependent on DOT1L. Upon *in vitro* activation, *Dot1L-*KO B cells failed to proliferate and after an *in vivo* immune challenge these cells failed to differentiate into GC B cells, and establish an effective immune response. In addition, we observed accelerated partial plasma cell differentiation in *Dot1L*-KO B cells *in vitro*. RNA-Seq data generated from naïve and *in-vitro* activated *Dot1L*-KO and WT B cells demonstrated a strong enrichment of genes associated with plasma cell differentiation among the genes up-regulated in *Dot1L* KO. However, *in-vitro* activated B cells failed to downregulate B220, indicating an incomplete differentiation of *Dot1L*-KO plasma cells^46^, which is in agreement with the failure to upregulate *Irf4*, a factor essential for plasma cell differentiation^50^. Our results also revealed that DOT1L supports MYC activity, which B cells depend on to differentiate effectively into pro-proliferative GC B cells^25^. Recent studies have shown that inhibition of DOT1L also leads to reduced *Myc* expression in multiple myeloma^68^ and in MYC-driven B cell lymphoma^69^, indicating that the connection between DOT1L and MYC has broad implications in B cell physiology. Furthermore, in neuroblastoma H3K79me2 methylation has been shown to be a strict prerequisite for MYC-induced transcriptional activation, indicating a mutual interaction between DOT1L and MYC^70^. Interestingly, this novel interaction has also been reported recently in colorectal cancer^71^. In addition to the crucial role in GC formation we also identify DOT1L as a critical factor in maintaining MZ B cells. Further exploring the strong reduction of MZ B cells upon loss of *Dot1L* should provide additional insights regarding the contribution of DOT1L in orchestrating normal B cell physiology.

In addition to supporting MYC and BACH2 activity, our findings also suggest that DOT1L supports the repression of target genes of PRC2. The observation that *Ezh2* is normally H3K79-dimethylated and downregulated in *Dot1L*-KO B cells, and that *Dot1L* and *Ezh2* are co-regulated in B cells indicates that DOT1L promotes repression of PRC2 targets by maintaining expression of *Ezh2*. A direct stimulatory effect of H3K79me on H3K27me3 synthesis is not likely to be involved since the modifications occur in distinct locations in the genome and are associated with opposite transcriptional states. Finally, besides MYC and PRC2, additional factors controlled by DOT1L may impact differentiation of B cells. For example, it will be interesting to determine the role of other candidate transcriptional regulators regulated by DOT1L (Tables 1-2). The number of transcriptional regulators involved in B cell development affected by DOT1L ablation implies the existence of a complex regulatory network that warrants further investigations to untangle.

Understanding the critical role of DOT1L in normal B cell differentiation is also relevant for disease. Considering the requirement for DOT1L in supporting humoral immune responses that we show here, targeting DOT1L may offer an opportunity for immune suppression. Given the strong dependency of GC B cells on DOT1L and the role of DOT1L in controlling the activity of EZH2 together with the oncogenic factor MYC, DOT1L inhibition may also offer a novel therapeutic angle in the treatment of diffuse large B cell lymphoma of the GCB type. In summary, in B cells, DOT1L has a central role in guiding dynamic epigenetic states controlling differentiation and ensuring functional immune responses, with a potential for clinical utility.

## Methods

### Mice

*Mb1-*Cre^*+/−*^*;Dot1L^fl/fl^* mice were derived by crossing the Dot1Ltm1a(KOMP)Wtsi line - generated by the Wellcome Trust Sanger Institute (WTSI) and obtained from the KOMP Repository (www.komp.org) - with the MB1-Cre strain kindly provided by M. Reth^72^. Mice from this newly created *Mb1-*Cre^*+/−*^;*Dot1L* strain were maintained under specific pathogen free (SPF) conditions at the animal laboratory facility of the Netherlands Cancer Institute (NKI; Amsterdam, Netherlands). Mice used for experiments were between 6-8 weeks old and of both genders. All experiments were approved by the Animal Ethics Committee of the NKI and performed in accordance with the Dutch Experiments on Animals Act and the Council of Europe.

### Genotyping PCR

Mice were genotyped for *Dot1L* using the forward primer (Dot1L: FWD, GCAAGCCTACAGCCTTCATC) and reverse primer 1 (Dot1L:REV1, CACCGGATAGTCTCAATAATCTCA) to identify WT allele; *Dot1L*^wt^ (642 Bps) while the floxed allele; *Dot1L*^fl^ (1017 Bps) was identified by using Dot1L: FWD and reverse primer 2 (Dot1L:REV2, GAACCACAGGATGCTTCAG). WT allele (418 Bps) for *Mb1* was detected by using forward primer (Mb1-FWD1: CTGCGGGTAGAAGGGGGTC) and reverse primer (Mb1-REV1: CCTTGCGAGGTCAGGGAGCC) while *Cre* (219 Bps) was detected by using forward primer (Mb1-FWD2: GTGCAAGCTGAACAACAGGA) and reverse primer (Mb1-REV2: AAGGAGAATGTGGATGCTGG).

### Flow cytometry

Single cell suspensions were made from bone marrow, spleen and Peyer’s patches. Bone marrow, spleen and blood samples were subjected to erythrocyte lysis. Distinct cellular populations were identified using combination of fluor-conjugated antibodies against surface markers (Table 3). Cells were stained with fluorescently labeled antibodies (Table 4). For intracellular staining cells were fixed and permeabilized using the Transcription Factor Buffer kit (Becton Dickinson, BD). Antibodies for intracellular staining were diluted in Perm/Wash buffer except for cleaved caspase-3. For H3K79me2 staining, cells were first stained with surface markers and fixed and permeabilized as described before. After fixation and permeabilization cells were washed with Perm/Wash containing 0.25% SDS. H3K79me2 specific antibody (Millipore) was diluted 1:200 into Perm/Wash + 0.25% SDS and cells were incubated for 30 min. Cells were washed with Perm/Wash and incubated with the secondary antibodies Donkey anti-Rabbit AF555 (Thermo Scientific) or Goat-anti-Rabbit AF488 (Invitrogen) 1:1000 in Perm/Wash. For cleaved caspase-3 staining the cells were first stained with surface markers and then fixed with 4% formaldehyde for 15 minutes at room temperature. After washing with 0.25% tween buffer the cells were permeabilized with 0.25% tween buffer for 15 minutes at room temperature. Following permeabilization the cells were stained with cleaved caspase-3 antibody as 1:50 diluted in 0.25% tween buffer for 45 minutes at room temperature under dark condition. Flow cytometry was performed using the LSR Fortessa (BD Biosciences) and data were analyzed with FlowJo software (Tree Star Inc.). Histograms were smoothed.

**Table 3:**
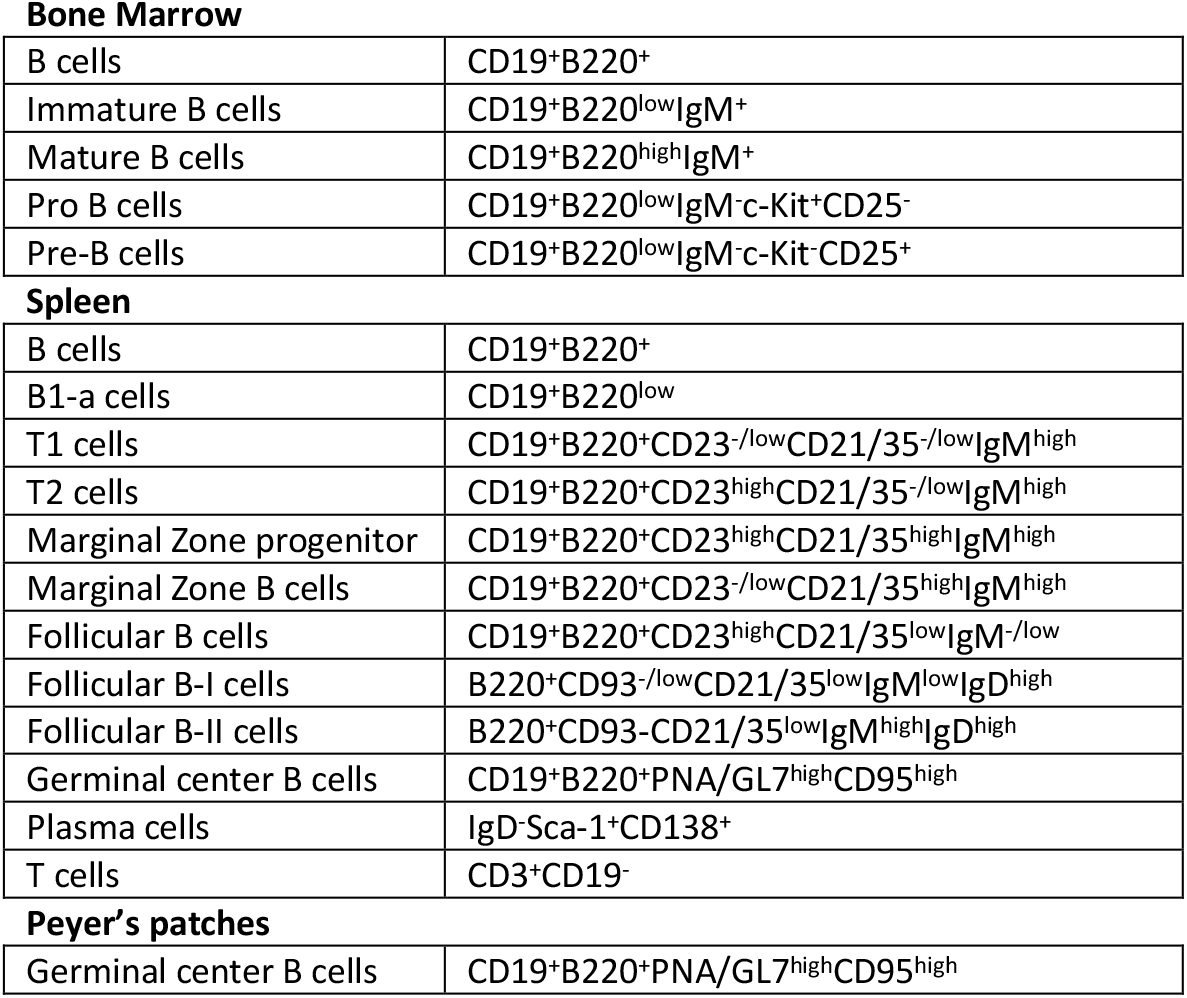
Surface markers for identification of specific cellular population.

**Bone Marrow**
|  |  |
| --- | --- |
| B cells | CD19<sup>+</sup>B220<sup>+</sup> |
| Immature B cells | CD19<sup>+</sup>B220<sup>low</sup>IgM<sup>+</sup> |
| Mature B cells | CD19<sup>+</sup>B220<sup>high</sup>IgM<sup>+</sup> |
| Pro B cells | CD19<sup>+</sup>B220<sup>low</sup>IgM<sup>-</sup>c-Kit<sup>+</sup>CD25<sup>-</sup> |
| Pre-B cells | CD19<sup>+</sup>B220<sup>low</sup>IgM<sup>-</sup>c-Kit<sup>-</sup>CD25<sup>+</sup> |
| **Spleen** |  |
| B cells | CD19<sup>+</sup>B220<sup>+</sup> |
| B1-a cells | CD19<sup>+</sup>B220<sup>low</sup> |
| T1 cells | CD19<sup>+</sup>B220<sup>+</sup>CD23<sup>-/low</sup>CD21/35<sup>-/low</sup>IgM<sup>high</sup> |
| T2 cells | CD19<sup>+</sup>B220<sup>+</sup>CD23<sup>high</sup>CD21/35<sup>-/low</sup>IgM<sup>high</sup> |
| Marginal Zone progenitor | CD19<sup>+</sup>B220<sup>+</sup>CD23<sup>high</sup>CD21/35<sup>high</sup>IgM<sup>high</sup> |
| Marginal Zone B cells | CD19<sup>+</sup>B220<sup>+</sup>CD23<sup>-/low</sup>CD21/35<sup>high</sup>IgM<sup>high</sup> |
| Follicular B cells | CD19<sup>+</sup>B220<sup>+</sup>CD23<sup>high</sup>CD21/35<sup>low</sup>IgM<sup>-/low</sup> |
| Follicular B-I cells | B220<sup>+</sup>CD93<sup>-/low</sup>CD21/35<sup>low</sup>IgM<sup>low</sup>IgD<sup>high</sup> |
| Follicular B-II cells | B220<sup>+</sup>CD93<sup>-</sup>CD21/35<sup>low</sup>IgM<sup>high</sup>IgD<sup>high</sup> |
| Germinal center B cells | CD19<sup>+</sup>B220<sup>+</sup>PNA/GL7<sup>high</sup>CD95<sup>high</sup> |
| Plasma cells | IgD<sup>-</sup>Sca-1<sup>+</sup>CD138<sup>+</sup> |
| T cells | CD3<sup>+</sup>CD19<sup>-</sup> |
| **Peyer's patches** |  |
| Germinal center B cells | CD19<sup>+</sup>B220<sup>+</sup>PNA/GL7<sup>high</sup>CD95<sup>high</sup> |

**Table 4:** Antibodies used for flowcytometry.

**Bone Marrow**
| Antibodies | Fluorophore | Clone | Dilution | Vendor |
| --- | --- | --- | --- | --- |
| CD19 | APC-H7 | 1D3 | 1:200 | BD Pharmingen |
| CD19 | PerCpCy5,5 | 1D3 | 1:200 | BD Pharmingen |
| CD45R (B220) | Pacific Blue | RA3-6B2 | 1:200 | BD Pharmingen |
| IgM | PECy7 | 11/41 | 1:200 | eBioscience |
| CD117 (cKit) | APC | 2B8 | 1:200 | eBioscience |
| CD25 | PE | PC61 | 1:200 | Biolegend |
| IgD | FITC | 11-26c.2a | 1:200 | BD Pharmingen |
| DAPI |  |  | 1:20 | Sigma-Aldrich |
| PI |  |  | 1:20 | Sigma-Aldrich |

**Spleen**
| Antibodies | Fluorophore | Clone | Dilution | Vendor |
| --- | --- | --- | --- | --- |
| GL-7 | FITC | GL7 | 1:200 | BD Pharmingen |
| Peanut Agglutinin | FITC |  | 1:400 | Vector |
| CD23 | BV421 | B3B4 | 1:200 | Biolegend |
| CD19 | PerCpCy5,5 | 1D3 | 1:200 | BD Pharmingen |
| CD19 | APC-H7 | 1D3 | 1:200 | BD Pharmingen |
| CD19 | A780 | 1D3 | 1:200 | eBioscience |
| CD95 | Biotin | 15A7 | 1:100 | eBioscience |
| CD95 | PE | Jo2 | 1:200 | BD Pharmingen |
| CD21/35 | APC | 7E9 | 1:200 | Biolegend |
| CD93 | BV650 | AA4.1 | 1:200 | BD Pharmingen |
| IgM | BV785 | II/41 | 1:200 | BD Pharmingen |
| IgD | BV650 | 11-26c.2a | 1:200 | Biolegend |
| IgD | PE | 11-26C | 1:800 | eBioscience |
| IgD | AF-488 | 11-26c2a | 1:200 | eBioscience |
| CD3 | FITC | 17A2 | 1:200 | Biolegend |
| CD3 | AF488 | 145-2C11 | 1:200 | eBioscience |
| CD45R (B220) | V500 | RA3-6B2 | 1:200 | BD Pharmingen |
| CD45R (B220) | PB | RA3-6B2 | 1:200 | BD Pharmingen |
| CD45R (B220) | APC | RA3-6B2 | 1:200 | BD Pharmingen |
| CD45R (B220) | BV510 | RA3-6B2 | 1:200 | Biolegend |
| CD138 | PE | 281-2 | 1:200 | BD Pharmingen |
| CD138 | BV605 | 281-2 | 1:200 | Biolegend |
| Streptavidin | BV605 |  | 1:400 | BD Pharmingen |
| Zombie NIR |  |  | 1:1000 | Biolegend |
| 7AAD |  |  | 1:500 | Biolegend |
| PI |  |  | 1:20 | Sigma-Aldrich |
| DAPI |  |  | 1:20 | Sigma-Aldrich |

**Peyer's Patches**
| Antibodies | Fluorophore | Clone | Dilution | Vendor |
| --- | --- | --- | --- | --- |
| CD45R (B220) | Pacific Blue | RA3-6B2 | 1:200 | BD Pharmingen |
| CD19 | PerCpCy5,5 | 1D3 | 1:200 | BD Pharmingen |
| IgM | APC | Polyclonal | 1:200 | Southern Biotech |
| Peanut Agglutinin | FITC |  | 1:400 | Vector |
| CD95 | PE | Jo2 | 1:200 | BD Pharmingen |
| DAPI |  |  | 1:20 | Sigma-Aldrich |

**Class switch recombination (*In vitro*)**
| Antibodies | Fluorophore | Clone | Dilution | Vendor |
| --- | --- | --- | --- | --- |
| CD19 | PerCpCy5,5 | 1D3 | 1:400 | BD Pharmingen |
| IgG1 | PE | 15H6 | 1:800 | Southern Biotech |
| IgM | APC | Polyclonal | 1:1000 | Southern Biotech |
| IgG3 | PE | Polyclonal | 1:800 | Southern Biotech |
| DAPI |  |  | 1:20 | Sigma-Aldrich |

**Plasma cells (*In vitro*)**
| Antibodies | Fluorophore | Clone | Dilution | Vendor |
| --- | --- | --- | --- | --- |
| CD138 | BV605 | 281-2 | 1:200 | Biolegend |
| CD45R (B220) | BV510 | RA3-6B2 | 1:200 | Biolegend |
| IgD | BV650 | 11-26c.2a | 1:200 | Biolegend |
| Sca1 | PeCy7 | D7 | 1:200 | Biolegend |
| CD3e | APC | 145-2C11 | 1:200 | BD Pharmingen |
| Ter119 | APC | TER-119 | 1:200 | BD Pharmingen |
| Blimp* | PE | SE7 | 1:200 | Biolegend |
| Zombie NIR |  |  | 1:1000 | Biolegend |
| * Intracellular |  |  |  |  |

**Cleaved Caspase-3 (*In vitro*)**
| Antibodies | Fluorophore | Clone | Dilution | Vendor |
| --- | --- | --- | --- | --- |
| CD19 | PerCpCy5,5 | 1D3 | 1:400 | BD Pharmingen |
| Cleaved Caspse-3 | AF488 |  | 1:50 | Cell signaling |
| Zombie NIR |  |  | 1:1000 | Biolegend |
| * Intracellular |  |  |  |  |

### Immunization

Adult mice were inoculated intravenously with sub-lethal dose 2×10^5^ PFU (Plaque forming units) of lymphocytic choriomeningitis virus strain Armstrong. Serum was collected prior to immunization and 14 days post immune challenge. For NP-CGG immunization mice were injected intraperitoneally with 100 μg of alumprecipitated NP-CGG [(4-hydroxy-3-nitrophenyl) acetyl coupled to chicken γ–globulin, BIOSEARCH™ TECHNOLOGIES] in a 200 μl of NP-CGG alum solution. To determine the serum titers of NP-specific IgM and IgG1, mice were bled from the tail vein on day 0,7,14,21,28, and 35.

### Class switch recombination and Proliferation

Single cells suspensions were prepared from the spleen of 6-8-week-old mice. Following erythrocyte lysis, naïve splenic B cells were enriched by the depletion of CD43 expressing cell using biotinylated anti-CD43 antibody (Clone S7, BD Biosciences), BD IMag Streptavidin Particles Plus and the IMag® system (BD Biosciences), as described by the manufacturer. To measure their proliferative capacity, naïve B cells were labeled for 10 min at 37°C with 5 μM Cell Trace Violet (CTV, Life Technologies, Invitrogen™) in IMDM medium containing 2% FCS,100 mM pen/strep and 100 mM β-mercaptoethanol. After washing, cells were cultured in complete IMDM medium (IMDM supplemented with 8% FCS, 100 mM pen/strep and 100 mM β-mercaptoethanol) at a density of 10^5^ cells/well in 24 well plates. CSR to IgG3 and IgG1 was induced in T cell independent manner by exposure to 5 μg/ml of Lipopolysaccharide (Escherichia Coli LPS, 055:B5, Sigma) or LPS+rIL-4. rIL4 was used at a concentration of 10 ng/ml. Cells were exposed to anti-CD40 (1ug/ml, BD Clone HM40-3) and rIL-4 (10 ng/ml) to induce IgG1 switching in a T cell dependent manner. Four days later, the cells were harvested and stained with CD19, IgM, and IgG3 (LPS cultures) or IgG1 (LPS/rIL4 cultures or anti-CD40/rIL-4) to determine CSR frequency along with CTV dilution as an indicator of cell multiplication.

### ELISA

LCMV-specific serum IgM and IgG levels were measured by ELISA. In short, Nunc-Immuno Maxisorp plates (Fisher Scientific) were coated overnight at 4°C with virus in bicarbonate buffer. Plates were subsequently incubated for 1h with blocking buffer (PBS/5% milk powder (Fluka Biochemika)). Sera from mice were diluted in PBS/1% milk powder and incubated for 1h at 37°C. Next, HRP-conjugated IgG and IgM antibodies (Southern Biotech) were diluted 1:4000 in PBS/1% milk powder and incubated 1h at 37°C. Plates were developed with TMB substrate (Sigma Aldrich), and the color reaction was stopped by the addition of 1 M H_2_SO_4_. Optical density was read at 450 nm (OD_450_) using a Microplate reader (Model 680, Bio-Rad). To quantify NP-specific serum antibodies, plates were coated with 2 μg/ml NP_30_-BSA. Serum was added at a starting dilution of 1:100 followed by 3-fold serial dilutions and incubated at room temperature for 2 hours. Bound serum antibodies were detected with polyclonal biotinylated goat anti-mouse IgM or anti-IgG1 (Southern Biotech), streptavidin-alkaline phosphatase conjugate (Roche) and chromogenic substrate 4-nitrophenyl phosphate (Sigma). Purified monoclonal antibodies (B1-8μ and 18-1-16y1) were used as standards for quantification.

### Lectin histochemistry

Lymphoid tissues such as spleens and lymph nodes were fixed in EAF (ethanol, acetic acid, formaldehyde, saline) for 24 hours and subsequently embedded in paraffin. 4 um-thick sections were stained with the lectin Peanut Agglutinin (PNA, Vector Laboratories) at 1:1500 dilution to reveal germinal centers. The sections were counterstained with hematoxyline.

### Sorting and *in vitro* activation for RNA and ChIP-Seq

For RNA sequencing, cells were first depleted for CD43+ cells and either subjected to MACS sorting for CD19+ cells as naïve B cells or activated for two days with LPS and IL-4. Following activation, the cells were enriched for CD19+ by MACS according to the manufacturer instructions. For ChIP-Seq, CD43-cells were either FACS sorted for CD19+ as naïve B cell pool or activated for three days with LPS and IL-4 and subjected to FACS sorting for CD19+ expression.

### RNA-Seq sample preparation

MACS sorted CD19+ cells were resuspended in Trizol (Ambion Life Technologies) and total RNA was extracted according to the manufacturer’s protocol. Quality and quantity of the total RNA was assessed by the 2100 Bioanalyzer using a Nano chip (Agilent). Only RNA samples having an RNA Integrity Number (RIN) > 8 were subjected to library generation.

### RNA-Seq library preparation

Strand-specific cDNA libraries were generated using the TruSeq Stranded mRNA sample preparation kit (Illumina) according to the manufacturer’s protocol. The libraries were analyzed for size and quantity of cDNAs on a 2100 Bioanalyzer using a DNA 7500 chip (Agilent), diluted and pooled in multiplex sequencing pools. The libraries were sequenced as 65 base single reads on a HiSeq2500 (Illumina).

### RNA-Seq preprocessing

Strand-specific RNA reads (11-33 million reads per sample), 65 bp single-end, were aligned against the mouse reference genome (Ensembl build 38) using Tophat (version 2.1, bowtie version 1.1). Tophat was supplied with a Gene Transfer File (GTF, Ensembl version 77) and was supplied with the following parameters: ‘--prefilter-multihits -no-coverage-search -bowtie1 -library-type fr-firststrand’. In order to count the number of reads per gene, a custom script which is based on the same ideas as HTSeq-count has been used. A list of the total number of uniquely mapped reads for each gene that is present in the provided Gene Transfer Format (GTF) file was generated. Genes that have no expression across all samples within the dataset were removed. Analysis was restricted to genes that have least 2 counts per million (CPM) value in all samples in specific contrasts, to exclude very low abundance genes. Differential expression analysis was performed in R language (version 3.5.1) on only relevant samples using edgeR package and default arguments with the design set to either Dot1LKO status, Ezh2KO status or cell type. Genes were considered to be differentially expressed when the False discovery rate (FDR) was below 0.05 after the Benjamini-Hochberg multiple testing correction. Sets of differentially expressed genes in indicated conditions were called ‘gene signatures. MA plots were generated after differential expression analysis carried by edgeR package^73, 74^. Read counts were corrected for gene length based on the longest transcript of the gene followed by normalization for the library size and shown as transcript per million (TPM). Counts are shown as counts per million after trimmed mean of M-values (TMM) normalization using the edgeR R package. For analyses where we performed expression matching, we chose genes with an absolute log2 fold changes less than 0.1 and false discovery rate corrected p-values above 0.05 that were closest in mean expression to each of the genes being matched without replacement. The RNA-Seq datasets reported in this article have been deposited at the National Center for Biotechnology Information under the accession number GSE138909.

### Ingenuity pathway analysis (IPA)

Lists of differentially expressed genes (FDR < 0.05) between WT and KO B cells both from naïve and activated conditions were submitted to IPA using default settings to identify potential upstream regulators.

### Gene set enrichment analysis (GSEA)

GSEA was carried out after differential expression analysis. Gene set enrichment was shown as barcode plot where the genes were ranked according to the log2 fold change between the compared conditions. Statistical significance for the enrichment of gene set was determined by Fast approximation to mroast (FRY) gene set test ^75^ from limma package and two-sided directional p value less than 0.05 was considered significant.

### Functional enrichment analysis

Functional enrichment analysis was carried by ‘g:GOST’ tool with default arguments from ‘g:Profiler2’ package and performed in R language (version 3.5.1).

### ChIP-Seq sample preparation

Sorted cells were centrifuged at 500 rcf. The pellet was resuspended in IMDM containing 2% FCS and formaldehyde (Sigma) was added to a final concentration of 1%. After 10 min incubation at RT glycine (final concentration 125 mM) was added and incubated for 5 min. Cells were washed twice with ice-cold PBS containing Complete, EDTA free, protein inhibitor cocktail (PIC) (Roche). Cross-linked cell pellets were stored at −80°C. Pellets were resuspended in cold Nuclei lysis buffer (50mM Tris-HCl pH 8.0, 10mM EDTA pH8.0, 1%SDS) + PIC and incubated for at least 10 min. Cells were sonicated with PICO to an average length of 200-500bp using 30 s on/ 30 s off for 3 min. After centrifugation at high speed debris was removed and 9x volume of ChIP dilution buffer (50mM Tris-HCl pH8, 0.167M NaCl, 1.1% Triton X-100, 0.11% sodium deoxycholate) + PIC and 5x volume of RIPA-150 (50mM Tris-HCl pH8, 0.15M NaCl, 1 mM EDTA pH8, 0.1% SDS, 1% Triton X-100, 0.1% sodium deoxycholate) + PIC was added. Shearing efficiency was confirmed by reverse crosslinking the chromatin and checking the size on agarose gel. Chromatin was pre-cleared by adding ProteinG Dynabeads (Life Technologies) and rotation for 1 hour at 4°C. After the beads were removed 2μl H3K79me1, 2 μl H3K79me2 (NL59, Merck Millipore) and 1 μl H3K4me3 (ab8580, Abcam) were added and incubated overnight at 4°C. ProteinG Dynabeads were added to the IP and incubated for 3 hours at 4°C. Beads with bound immune complexes were subsequently washed with RIPA-150, 2 times RIPA-500 (50mM Tris-HCl pH8, 0.5M NaCl, 1mM EDTA pH8, 0.1% SDS, 1% Triton X-100, 0.1% sodium deoxycholate), 2 times RIPA-LiCl (50mM Tris-HCl pH8, 1mM EDTA pH8, 1% Nonidet P-40, 0.7% sodium deoxycholate, 0.5M LiCl2) and TE. Beads were resuspended in 150 μl Direct elution buffer (10mM Tris-HCl pH8, 0.3M NaCl, 5mM EDTA pH8, 0.5%SDS) and incubated overnight at 65°C and input samples were included. Supernatant was transferred to a new tube and 1 μl RNase A (Sigma) and 3 μl ProtK (Sigma) were added per sample and incubated at 55°C for 1 hour. DNA was purified using Qiagen purification columns.

### ChIP-Seq Library preparation

Library preparation was done using KAPA LTP Library preparation kit using the manufacturer’s protocol with slight modifications. Briefly, after end-repair and A-tailing adaptor were ligated followed by Solid Phase Reversible Immobilization (SPRI) clean-up. Libraries were amplified by PCR and fragments between 250-450 bp were selected using AMPure XP beads (Beckman Coulter). The libraries were analyzed for size and quantity of DNAs on a 2100 Bioanalyzer using a High Sensitivity DNA kit (Agilent), diluted and pooled in multiplex sequencing pools. The libraries were sequenced as 65 base single reads on a HiSeq2500 (Illumina).

### ChIP-Seq preprocessing

ChIP-Seq samples were mapped to mm10 (Ensembl GRCm38) using BWA-MEM with the option ‘-M’. Duplicate reads were removed using MarkDuplicates from the Picard toolset with ‘VALIDATION_STRINGENCY=LENIENT’ and ‘REMOVE_DUPLICATES=true’ as arguments. Bigwig tracks were generated from these bam files by using bamCoverage from deepTools using the following arguments: ‘-of bigwig -binsize 25 -normalizeUsing RPGC - ignoreForNormalization chrM -effectiveGenomeSize 2652783500’. Bigwig files were loaded into R using the ‘import.bw()’ function from the rtracklayer R package for visualization of heatmaps and genomic tracks. TSSs for heatmaps were taken from Ensembl GRCm38.77 gene models by taking the first base pair of the 5’ UTR of transcripts. When such annotation was missing, the most 5’ position of the first exon was taken. The ChIP-Seq datasets reported in this article have been deposited at the National Center for Biotechnology Information under the accession number GSE138906.

### Statistics

Statistical analyses were performed using Prism 7 (GraphPad). Data are presented as mean ±SD except for Fig. 4D where it is presented as mean± SEM. The unpaired Student’s t-test with two-tailed distributions was used to calculate the p-value. A p-value < 0.05 was considered statistically significant.

## Acknowledgements

We like to thank the core facilities of the NKI-AVL for their superb biotechnical and molecular help. We highly appreciate the support from Michael Reth in providing his *Mb1*-Cre mouse model system.

## Author contributions

Conception and design, M.A.A, M.F.A., E.M.K-M., F.v.L., and H.J.

Acquisition of data, M.A.A., M.F.A., E.M.K-M., M.C., I.N.P., T.v.W., J-Y.S., and F.I.M.

Analysis and interpretation of data, M.A.A, M.F.A., E.M.K-M., R.A., E.d.W., K.R., F.v.L., and H.J.

Bioinformatics Analysis, M.A.A, T. v.d. B., M.F.A, E.d.W., and I.d.R.

Writing of the manuscript, M.A.A., M.F.A., E.M.K-M., F.v.L. and H.J.

